# DNA-protein interaction is dominated by short anchoring elements

**DOI:** 10.1101/2023.12.11.571177

**Authors:** Hong Chen, Yongping Xu, Hao Ge, Xiao-dong Su

**Affiliations:** State Key Laboratory of Protein and Plant Gene Research, School of Life Sciences, and Biomedical Pioneering Innovation Center (BIOPIC), Peking University, Beijing, 100871, China; Beijing International Center for Mathematical Research (BICMR) and Biomedical Pioneering Innovation Center (BIOPIC), Peking University, Beijing, 100871, China

**Author notes:** To whom correspondence should be addressed. Xiao-dong Su.

## Abstract

To understand the regulation of gene expression, it is essential to elucidate the binding mechanism of DNA binding domain (DBD) of transcription factors (TFs), and predict the location of transcription factor binding sites (TFBSs). For an exhaustive search of TFBSs, we have investigated four typical TFs with diverse origins, such as WRKY, PU.1, GLUCOCORTICOID RECEPTOR (GR), and MYC2 by using a newly developed method, KaScape. During KaScape experiments, we identified short sequences (3-4 bases) or “anchoring element” (AE) for the four TFs that dominated the bound population of DNA-DBD binding. We further developed the AEEscape (AE Energy landscape) algorithm to detect and confirm the AE and derived its binding energy landscape for all possible sequences. Our analysis of the energy landscape revealed an energetic funnel around the TFBS, which is related to the AE density gradient in the region surrounding the TFBS. Our results provide novel insights into the mechanism of TF binding to TFBSs.

## INTRODUCTION

Gene expression is essential for cell development and is primarily regulated at the transcriptional level (1). Transcription factors (TFs) can bind to their target sites for a period of time to recruit other factors to promote or prevent binding of the transcriptional machinery, therefore to control gene expression (2). However, it is challenging to identify DNA sequences that can be bound by specific TFs to precisely regulate gene expression (3). The double stranded DNA (dsDNA) segments that are bound by TFs in the cell are called transcription factor binding sites (TFBSs). The sequence required for TF binding is often referred to as a DNA binding motif. And the motif is usually characterized as a TFBS consensus by position weight matrix (PWM) (3-5). A PWM provides a score for all possible bases at each position in a binding site according to the position frequency matrix (PFM) (6)

There are many experimental methods to find TFBS motifs, including in vitro methods such as HT-SELEX (7) (high-throughput systematic evolution of ligands by exponential enrichment), PBM (8) (protein binding microarray), MITOMI (9,10) (mechanically induced trapping of molecular interactions), and recently KaScape (11), and in vivo methods such as ChIP-chip, ChIP-seq (chromatin immunoprecipitation followed by high-throughput sequencing) and their derived methods (12). Computational methods have also been developed in the recent years (3,13-16). Current computational methods for DNA binding site analyses fall into two categories. One is to take a collection of known binding sites and generate a representation to aid in the search and prediction of additional binding sites. The other one is to discover the location of the TFBSs in the DNA sequence (14). The sequence patterns which are degenerate DNA sequences containing A, C, G, T, and degenerate IUPAC nucleic acid codes, and position weight matrices (PWMs) are widely used as a model to describe TFBSs. Compared to sequence patterns, PWM can quantitatively characterize DNA binding sites with the position-independent assumption. A good PWM model relies on accurate sequence alignments, which is usually challenging (17,18). Because the TFBSs are in general degenerative, many sequences may match a given motif in the genome when the motif is used to scan for binding sites, leading to false positives. However, if we use the highest information consensus (i.e. high affinity binding sites) to scan, we could miss the weaker yet still functional binding sites (16). To solve this problem, higher-order models that take into account the interdependence between positions have been developed (19-22). Most of them used dinucleotide interactions information and did not consider the structural information of the DNA-TF complex (19,20).

The function of TFs is highly dependent on their fast and accurate recognition ability (23). One mechanism proposed to achieve the fast search ability of TFs is “Facilitated Diffusion” (FD), proposed by Berg and Von Hippel to solve the paradox (24) that the rate of TF finding its target (i.e. the association rate of *Escherichia coli* Lac repressor) greatly exceeds the diffusion-limited rate (25-27). The FD hypothesis combines one-dimensional diffusion along the DNA with three-dimensional diffusion near dsDNA. The FD mechanism implies that the region flanking the TFBSs influences TFs to search and find their targets. A binding energy funnel of RNA polymerase (RNAP) has been revealed (28). And the paper shows that the energetic funnel is related to the AT gradients around the RNAP binding site and can increase the probability of target finding. Besides AT gradients, repeats of degenerate recognition sites are reported in regions flanking the eukaryotic genes (29-31). And it is classified as gene regulatory regions (RR) (32). According to the research on the DNA binding domain of a TF, engrailed, RR can significantly enhance the binding affinity, which can ensure the local supply of TF (33). Although RR can trap TF, which slows down FD, a Brownian dynamics simulation shows that organizing binding energies into a funnel (appropriately positioning traps) can reduce the search time (34).

A recent research reported that short tandem repeats (STRs) can bind with TFs to tune eukaryotic gene expression. They anticipated that low-affinity binding sites can increase binding and proposed the STRs function as “rheostats” to tune local TF concentration and binding responses (35). Another research pointed out that the molecular dynamics-derived trinucleotide and tetranucleotide-based structure and energy parameters exist in a signature profile at the transcription start sites region of both prokaryotes and eukaryotes (36). However, the mechanism by which eukaryotic TFs find their target through flanking DNA has not been characterized, experimentally observed, or demonstrated at the molecular level.

In this paper, we describe a new concept named “anchoring element” (AE) which includes short segments (3-4 bp) essential for TFs to recognize and bind their binding sites. We have firstly discovered the existence of AEs during KaScape experiments (11), and demonstrated that the AE is a useful concept for understanding the mechanism by which TFs recognize and bind to dsDNA. We further developed an energy calculation algorithm AEEscape (AE Energy landscape) based on BEESEM (Binding Energy Estimation on SELEX with Expectation Maximization) algorithm (37) to analyze our experimental data. The BEESEM algorithm builds on a comprehensive biophysical model of protein-DNA interactions and is trained using the expectation-maximization method (37). The energy term used in BEESEM is the energy of the short DNA sequence physically covered by the protein and is represented by the position weight matrix parameters, which assumes that the bases are independent, whereas in the AEEscape algorithm, the energy term is the energy of the shortest sequence that can be specifically distinguished by the protein, and the short sequence at different locations of the random bases has been directly set as parameters. The AEEscape result shows that when the short sequence length is the same as the AE, its patterns are similar and the AE is the same with the lowest binding energy at different locations of the random sequence, when the short sequence length is shorter, the patterns are different, when the short sequence length is longer than the AE, almost all the sequences containing AE have low binding energy. This allows us to see that the AE dominates the binding energy of protein and DNA. Using the energy landscape generated by the AEEscape, we revealed an energetic funnel around the TFBS. We also find that the AE density gradient in the energy funnel, namely, the AE density around the TFBS, is important for the TF to find its target quickly and reliably.

## MATERIAL AND METHODS

### Experimental data generated from the KaScape method

The KaScape experiments (previously described (11)) were run for the TFs (the N-terminal DNA binding domain (DBD) of *At*WRKY1 (*At*WRKY1N), PU.1 DBD, the N-terminal DBD of cGAS (cGASN), GLUCOCORTICOID RECEPTOR (GR) DBD, and MYC2 DBD). Each KaScape experiment generates input and output sequencing reads (input-output dataset). The input data are random sequences before interacting with TF (the random dsDNA pool sequencing data) and the output data are sequences interacting with TF (the bound dsDNAs sequencing data). The random sequence types used in the KaScape experiments are shown in Supplementary Text 1. The random sequence types R1N4, R1N5, R1N6, and R1N7 were used to interact with *At*WRKY1N in each KaScape experiment, which are the same data sets as in the paper (11). For other TFs in each KaScape experiment, random sequence type R2N7 interacted with PU.1 DBD, cGASN, and GR DBD, respectively, while random sequence type R1N7 interacted with MYC2 DBD.

We have modified the KaScape experiment. In the modified KaScape protocol, we reduced the reaction system by a factor of 10 while maintaining the same molar concentration, omitted the wash step, collected the unbound dsDNAs, and sequenced the unbound dsDNAs (unbound data). Each modified KaScape generates unbound and output sequencing reads (unbound-output dataset). Random sequence type R1N5 was used to interact with *At*WRKY1N using the modified KaScape protocol (see Table S1).

In the results section, we mostly used the input-output dataset. If an unbound-output dataset was used, it would be indicated.

### Biophysical model of AEEscape

The biophysical model of AEEscape characterizes the binding energy of each m-mer element at different locations in the random base region. It is based on the BEESEM model (37). In the BEESEM model, the binding ability of a specific m-mer sequence ***s***_***j***_ is characterized by calculating the ratio (denoted by ***R***_***j***_) of ***P***(***s***_***j***_|***B***) (the fraction of TFBSs where the binding sequence is ***s***_***j***_) and 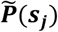 (the fraction of ***s***_***j***_ before binding to the TF) (Equation 1). ***P***(***B***) is the binding probability of all types of dsDNA sequences. ***u*** is the chemical potential of the TF. ***E***(***s***_***j***_) is the binding energy of ***s***_***j***_ regardless of its location on any sequence and is calculated by PWM. PWM assumes that the bases in the motif are independent of each other (38). Changing one base in the motif does not affect the energy contribution of the nearby unchanged base. However, there are correlations between bases when bound by TF, which can provide insight into the binding mechanism (39,40). To investigate the binding mechanism, we developed the AEEscape algorithm. In AEEscape, we assume that the binding site length m of ***s***_***j***_ is relatively short (2, 3, or 4). Due to the influence of the flanking fixed sequence, the binding energy landscape of ***s***_***j***_ may be different at different locations in the random bases. Therefore, 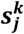 was introduced to represent a specific m-mer sequence at the kth location in the random bases. For simplicity, we only consider TFs that are bound to random base regions. We denote the length of random bases in each sequence by l. There are l-m+1 binding sites. 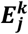 is the energy of 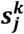. We have generalized Equation 1 by replacing ***s***_***j***_ with 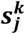 (Equation 2). Instead of using PWM to represent the energy as BEESEM did, in AEEscape we take 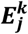 directly as a parameter. Finally, we use the EM algorithm to compute 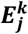, the chemical potential, and the auxiliary parameters. The loss function is shown in Equation 3 (see Supplementary Text 2 for a detailed description).

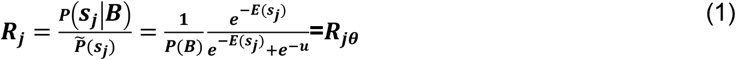

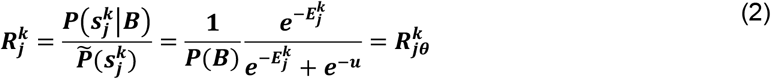

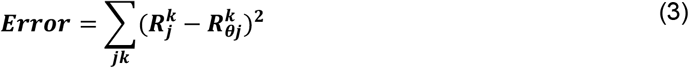

### Training and Evaluation of AEEscape

There are two types of KaScape data sets that are used to train the AEEscape model (described above). They are the input dsDNA (***P***(***S***_***i***_)) and the output dsDNA (***P***(***S***_***i***_|***B***)) (input-output dataset) as well as the unbound dsDNA (***P***(***S***_***i***_|***U***)) and the output dsDNA (unbound-output dataset). For the second data set, we use ***P***(***S***_***i***_|***U***) and ***F***_***UB***_ (the ratio of total sequencing reads in unbound versus bound) to represent the ratio of ***P***(***S***_***i***_) and ***P***(***B***) (Equation S15) and rearrange Equation S9 to obtain 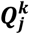 (Equation S16), which is similar to 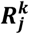. The loss function is shown in Equation 4. (see Supplementary Text 2 for a detailed description).

The binding energy for each sequence ***S***_***i***_ is denoted as ***E***(***S***_***i***_) and is calculated as Equation 5. We can evaluate the training effects by comparing the binding energy of sequences obtained directly from the experimental dataset with those predicted by the model parameters.

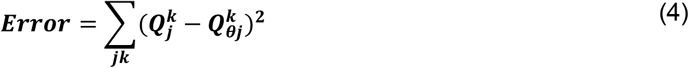

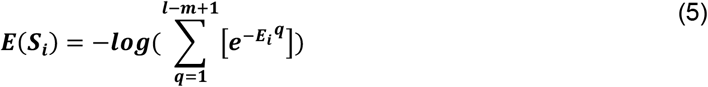

## RESULTS

### The discovery of the AE

To study the interaction mechanism between the protein and dsDNA, we analyzed the relative binding energy landscapes from a series of KaScape experiments for *At*WRKY1N in the KaScape paper (11), shown in Fig.1 A1 - Fig.1 A4. With the color changing from dark purple to yellow, the relative binding energy is getting smaller. As the high-affinity signals are less than 0, the relative binding energies bigger than 0 are shown in dark purple to conveniently observe the signals. When the random sequence type is R1N4, the relative binding energies of sequences containing GAC (sequences in white boxes, NGAC, or GACN) or GTC (sequences in red boxes, NGTC, or GTCN) are the lowest or relatively low. The GAC and GTC are in reverse complementary. The relative binding energies of all 16 sequences that contain GAC or GTC (annotated in the white or red boxes) are below 0 which suggests the *At*WRKY1N can specifically recognize the 3-mer GAC (or GTC in reverse complementary). When the random sequence type is R1N5, R1N6, or R1N7, the relative binding energies in the upper left square are the lowest or relatively low where the sequences are GACNN, GACNNN, and GACNNNN respectively with a slightly more noisy background. To test whether the *At*WRKY1N can specifically recognize the 3-mer GAC (or GTC in reverse complementary), we did a KaScape experiment for *At*WRKY1N with R1N3 random sequence, see Fig.1 B1. The lowest relative binding energy is GTC and GAC annotated in red. To reduce experimental noise, we modified the KaScape experiment to sequence both the bound and unbound random sequences. The relative binding energy landscape of the KaScape experiment for *At*WRKY1N and R1N5 is shown in Fig.1 B2. The sequences in the white box and the red box contain GAC and GTC respectively. From Fig.1 B2 we can see a fractal-like pattern. The fractal-like pattern has been revealed before when the authors used the K-mer graph to visualize the genomes of organisms (41). In their paper, it is suggested that the fractal-like pattern is caused by the under-representation of those strings that contain a particular short sequence of nucleotides. Since the relative binding energy shows a decrease as the color shifts from purple to yellow, the fractal-like pattern found here is due to those sequences with low relative binding energy values that contain a certain short sequence of nucleotides. For *At*WRKY1N, the fractal-like pattern is due to those sequences with low relative binding energy that contain the same short sequence of nucleotides GAC or GTC (in reverse complementarity). The relative binding energies of the majority of the sequences that contain GAC or GTC are below 0 (73 sequences’ relative binding energies are below 0 with the total 96 sequences) and most of the sequences’ relative binding energies are the lowest. The results shown here further suggest GAC (or GTC in its reverse complementary) is specific for *At*WRKY1N to bind and dominate the binding of *At*WRKY1N with dsDNA. The dominating short sequence GAC (or GTC) is defined as AE.

**Figure 1.**
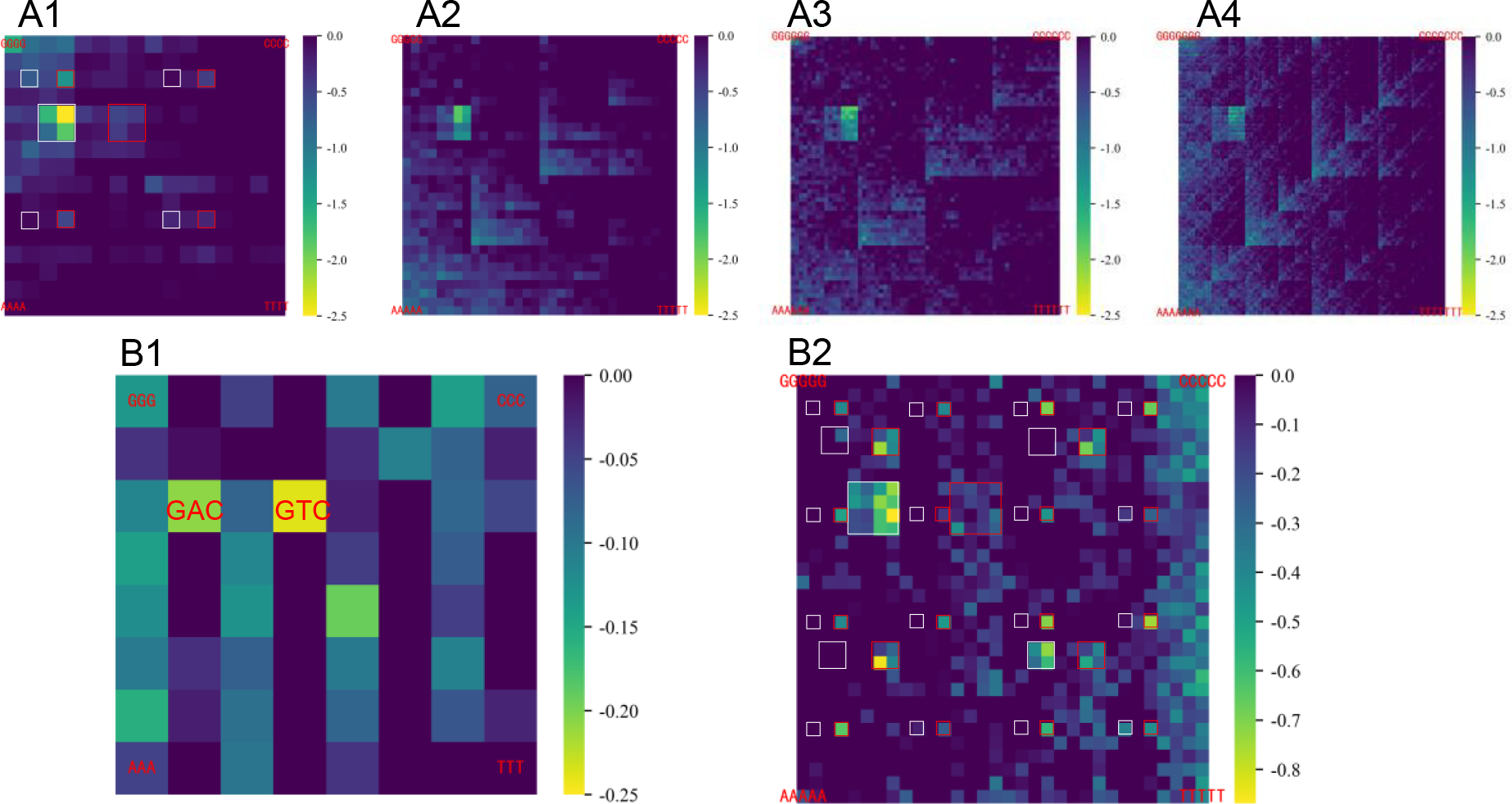
The KaScape binding energy landscapes on K-mer graphs for *At*WRKY1N with different random sequence types used in KaScape experiments. The relative binding energies bigger than 0 are shown in dark purple. (A1—A4) The KaScape experiments are the same in the paper (11). The random sequence type is R1N4, R1N5, R1N6, and R1N7 respectively. (B1) The random sequence type is R1N3. The sequences of low energy are labeled in red. (B2) The random sequence type is R1N5. The dataset used here is the unbound-output dataset. The sequences in the white boxes are GACNN, NGACN, or NNGAC. The sequences in the red boxes are GTCNN, NGTCN, or NNGTC. N represents G, C, A, or T.

To identify and confirm the existence of the AE, as well as to explore the binding energy of the AE and compare it with the other possible short sequences, we developed AEEscape, a tool that can be used to calculate the binding energy for all types of m-mer (called an element) short sequences at any location in the random base region. Figure 2 shows the predicted binding energies for m-mer at each location in the random base region for *At*WRKY1N. When m is set to 2 (Fig. 2 A1 – Fig. 2 A4), the binding energy landscape maps are different at each location (see also Fig. S1). When the value of m is set to 3 (Fig.2 B1 – Fig.2 B3), the landscape maps of the binding energy show a high degree of similarity. Specifically, the 3-mer sequence with the lowest binding energy at each location within the random base region is GAC (or its reverse complementary sequence, GTC) (Fig. S2). This implies that *At*WRKY1N is able to recognize the short 3-mer sequence GAC (or GTC) with specificity, regardless of its binding site. When m is equal to 4 (Fig. 2 C1 and Fig. 2 C2), the 4-mer sequences with the lowest binding energies at the first location in the random base region are GACT, GACC, and TGAC (Fig. S3A), while at the second location in the random base region, the 4-mer sequences with the lowest binding energies are GGTC, AGTC, CGTC, and GTCA (Fig. S3B). The 4-mer sequences with the lowest binding energies all contain GAC (or its reverse complementary sequence, GTC). By comprehensively examining the binding energies for 2-mers, 3-mers, and 4-mers at different sites within the random base region, it was determined that a 3-mer sequence is the shortest possible sequence that contains the necessary information for *At*WRKY1N to interact with dsDNA in a specific manner and dominate the binding energy of *At*WRKY1N interacting with dsDNA. To assess the accuracy and interpretability of the predicted result, we performed a direct comparison between the binding energy obtained from the experimental data (Fig. S4A) and the binding energy predicted by AEEscape (Fig. S4B). The pattern of the two figures shows a remarkable similarity. Our analysis revealed a correlation coefficient of 0.911 (Fig. S4C), providing strong evidence for the high degree of convergence, accuracy, and interpretability of the prediction. Therefore, our results have led us to conclude and validate that the AE (GAC or its reverse complementary sequence, GTC) serves as the most concise and accurate sequence for WRKY binding to dsDNA. In addition, our investigations have demonstrated that the binding energy of the AE is the lowest among all other potential m-mer sequences. However, the binding energy of the AE is only moderately lower than that of other possible m-mer sequences.

**Figure 2.**
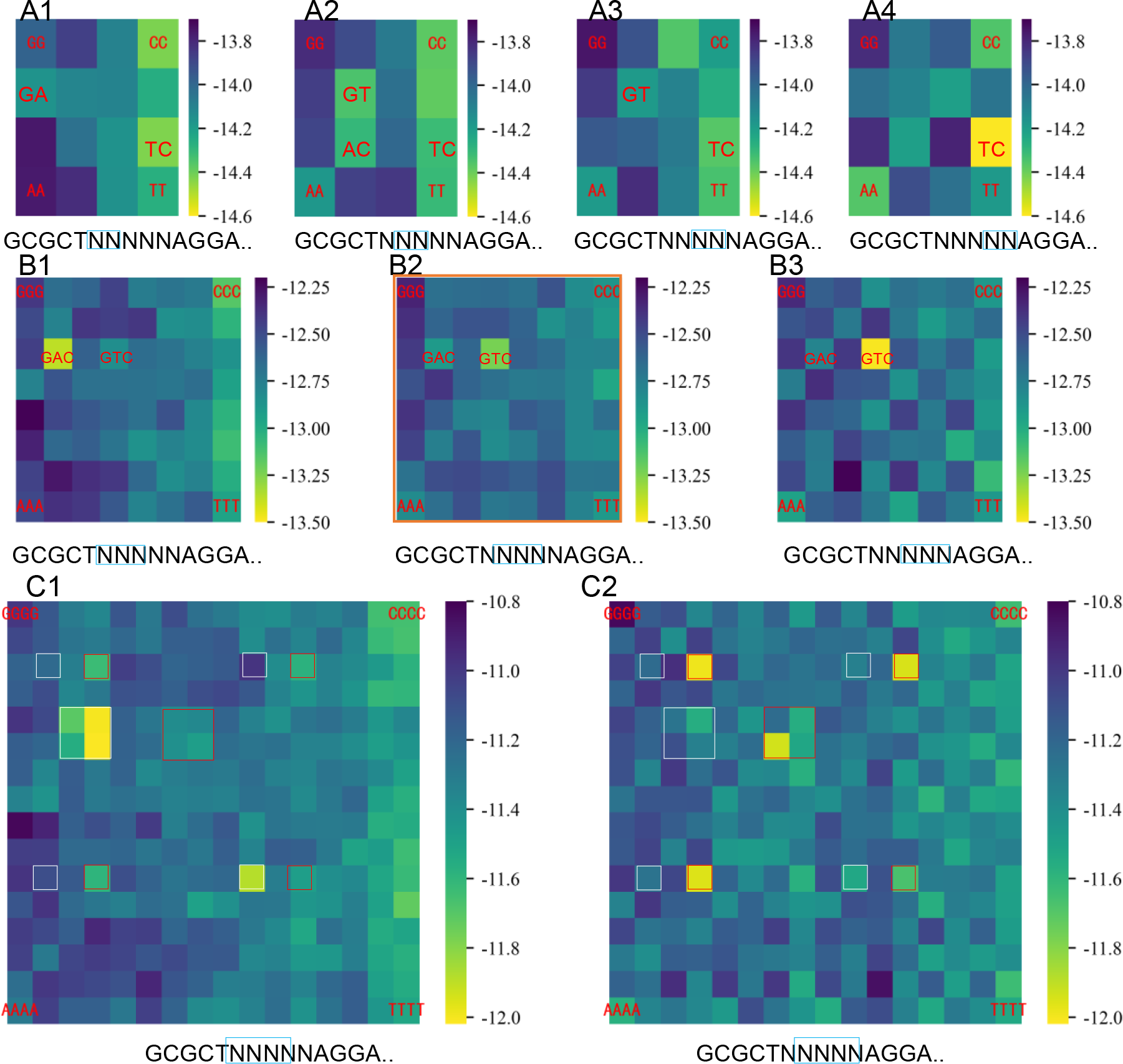
The binding energy landscapes at different locations in the random base region predicted by the AEEscape algorithm trained from KaScape experimental data for *At*WRKY1N. (The experimental data set is the same as the data used in Fig. 1B2). The binding energy parameters in each row are simultaneously predicted by training on one type of KaScape experiment. The sequences of low energy are labeled in red. The random sequence at the bottom of each figure is the sequence type used in the KaScape experiment. The blue box in the random region indicates the location of the predicted binding energy landscape. The binding energy in the orange box is used to calculate the binding energy near the TFBS. N represents G, C, A, or T. (A1 – A4). The base length in the model is 2. The location in the random base region is 1, 2, 3, and 4, respectively. (B1 – B3) The base length in the model is 3. The location in the random base region is 1, 2, and 3, respectively. (C1 – C2) The base length in the model is 4. The location in the random base region is 1 and 2, respectively. The sequences in the white boxes are GACN or NGAC. The sequences in the red boxes are GTCN or NGTC.

### The AE evidence of other DBDs

To explore whether the AE is ubiquitous, we performed the KaScape experiment (11) for other DNA binding proteins such as PU.1 DBD, cGASN, GR DBD, and MYC2 DBD. Figure 3 shows the relative binding energy landscape maps based on the K-mer graph for these DNA binding domains. On each map, we also found a fractal-like pattern. This suggests AE exist in these DBDs. For PU.1 DBD, the fractal-like pattern is due to those sequences with low relative binding energy containing the AE (GGAA or TTCC in reverse complementarity) (Fig. S5A). For cGASN, the fractal-like pattern is due to those sequences with low relative binding energy containing the AE (CAC) (Fig. S5B). As for GR DBD and MYC2 DBD, the background noise is larger, however, AEEscape can help us find the AEs for them. Figure 4 shows the predicted landscape of binding energy at each location in the random base region by AEEscape. Each row represents one protein type. From top to bottom, they are PU.1 DBD, cGASN, GR DBD, and MYC2 DBD. For PU.1 DBD, the relative binding energy of GGAA (or TTCC in reverse complementarity) is the lowest or the second lowest at all locations in the random base region (Fig.4 A1—Fig.4 A4 and Fig. S6). The AE for PU.1 is GGAA (or TTCC in reverse complementarity). For cGASN, the relative binding energy of CAC is the lowest at all locations in the random base region ((Fig.4 B1—Fig.4 B5 and Fig. S7). The AE of cGASN is CAC. For GR DBD, the relative binding energy of TGT (or ACA in reverse complementarity) is the lowest or relatively low at all locations in the random base region (Fig.4 C1—Fig.4 C5 and Fig. S8). The AE of GR DBD is ACA (or TGT in reverse complementarity). For MYC2 DBD, the relative binding energy of CAC (or GTG in reverse complementarity) is the lowest or relatively low at all locations in the random base region (Fig.4 D1—Fig.4 D5 and Fig. S9). The AE of MYC2 DBD is CAC (or GTG in reverse complementarity). Figure S10 shows a comparison of the binding energy profiles obtained from experimental measurements and those predicted by AEEscape. The close agreement between the predicted and experimental binding energy maps provides compelling evidence for the accuracy and interpretability of the prediction. In addition, the AEEscape algorithm is effective in filtering noise from the data.

**Figure 3.**
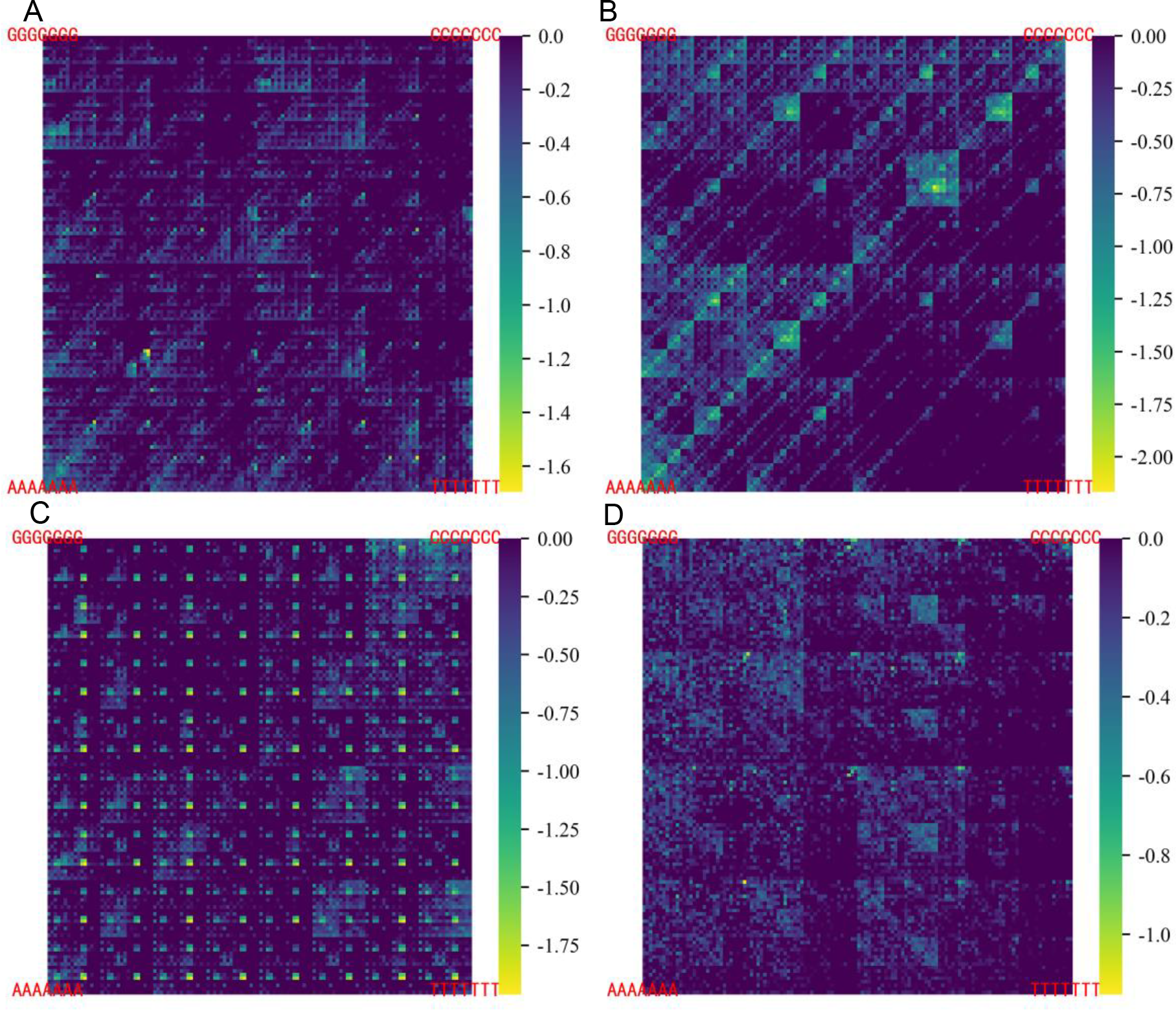
The KaScape binding energy landscape on a K-mer graph for different DNA binding domains of TFs. The relative binding energies bigger than 0 are shown in dark purple. (A) The KaScape experiment for PU.1 DBD. The random sequence type is R2N7. (B) The KaScape experiment for the cGASN. The random sequence type is R2N7. (C) The KaScape experiment for the GR DBD. The random sequence type is R2N7. (D) The KaScape experiment for the MYC2 DBD. The random sequence type is R1N7.

**Figure 4.**
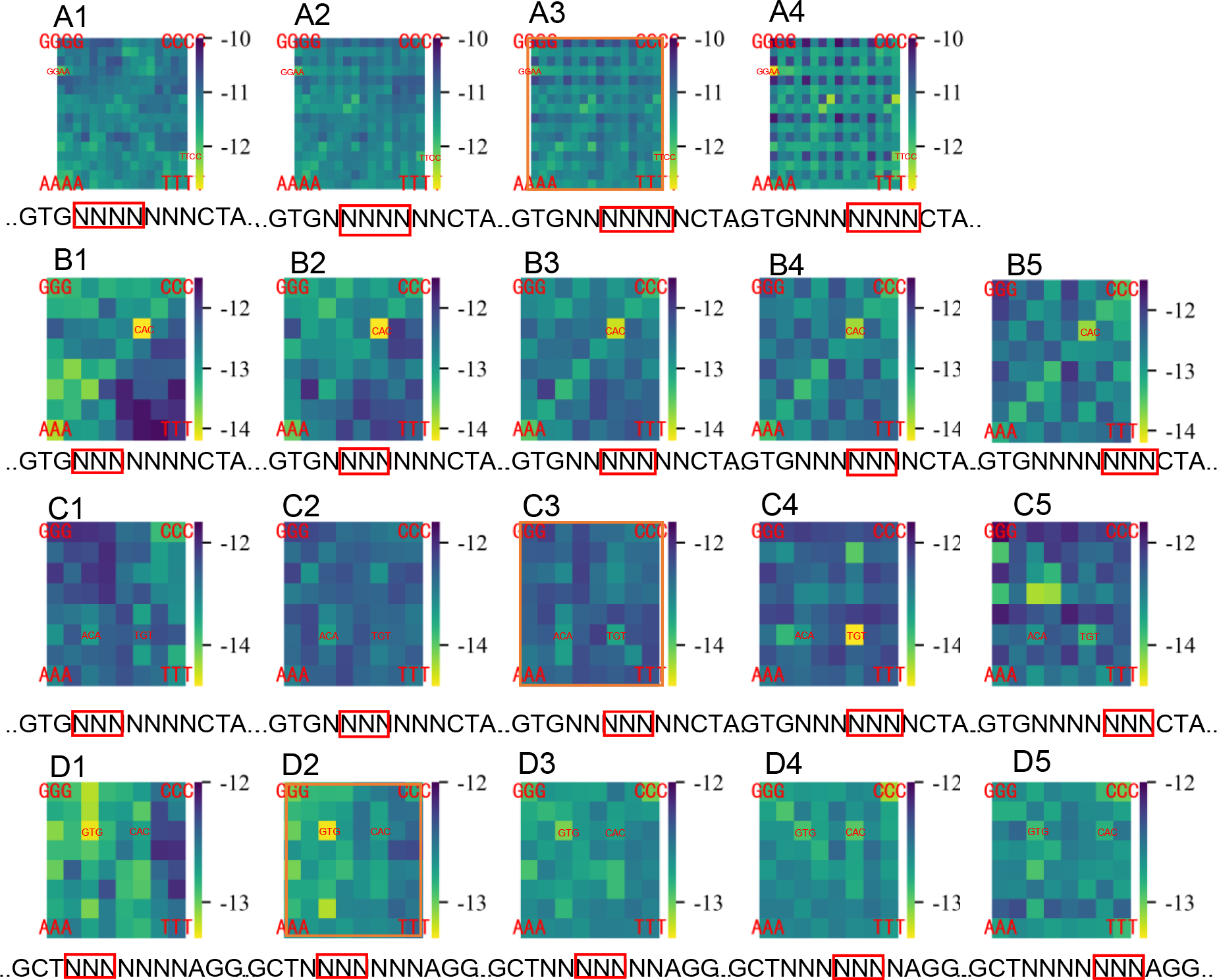
The binding energy landscapes at different locations in the random base region predicted by the AEEscape algorithm trained from KaScape experimental data for different proteins. The binding energy parameters in each row are simultaneously predicted from one type of KaScape experiment for each protein. The sequences of low energy are labeled in red. The random sequence at the bottom of each figure is the sequence type used in the KaScape experiment. The blue box in the random region indicates the location of the predicted binding energy landscape in the random region. The binding energy in the orange box is used to calculate the binding energy near the TFBS. (A1 – A4) The protein of KaScape experimental data used in training is PU.1 DBD (The experimental data is the same as the data used in Fig. 3A). The base length in the model is 4. The location in the random base region is 1, 2, 3, and 4, respectively. (B1 – B5) The protein of KaScape experimental data used in training is cGASN (the experimental data is the same as the data used in Fig. 3B). The base length in the model is 3. The location in the random base region is 1, 2, 3, 4, and 5, respectively. (C1 – C5) The protein of KaScape experimental data used in training is GR DBD (the experimental data is the same as the data used in Fig. 3C). The base length in the model is 3. The location in the random base region is 1, 2, 3, 4, and 5, respectively. (D1 – D5) The protein from the KaScape experimental data used in training is MYC2 DBD. (the experimental data is the same as the data used in Fig. 3D). The base length in the model is 3. The location in the random base region is 1, 2, 3, 4, and 5, respectively.

### AE gradient forming the energetic funnel

By far, our study has determined the binding energy landscape of the shortest m-mer sequences that contribute most to the specific interaction between TFs and double-stranded DNA. We then investigated the binding energy around TFBS annotated in the ChIP-seq dataset for WRKY (42), PU.1 (GSM1010843) (43), GR (44), and MYC2 (45), to gain insight into the process of TFBS search. We extracted the sequences from the genome of TFBS indicated by ChIP-seq data and filtered those sequences containing N. (see Table S2 for the ChIP-seq sequence numbers used). Then, we calculated the normalized binding energy (a function of the distance r from the target site 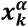 (the kth target site of the ***α***th TF)) adapted from the paper (28) (Equation 6). **⟨*Y*⟩** is the average of Y total target sequences (Equation 7). ***N***_***TF***_ is the number of TFs and ***M***_***α***_ is the number of target sequences for ***α***th TF. The energies framed by an orange box in Fig. 2 and Fig. 4 are used to infer 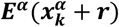 at each location of the target sequences. The function E(r) shows a funnel around the TFBS for those four TFs (Fig. 5).

**Figure 5.**
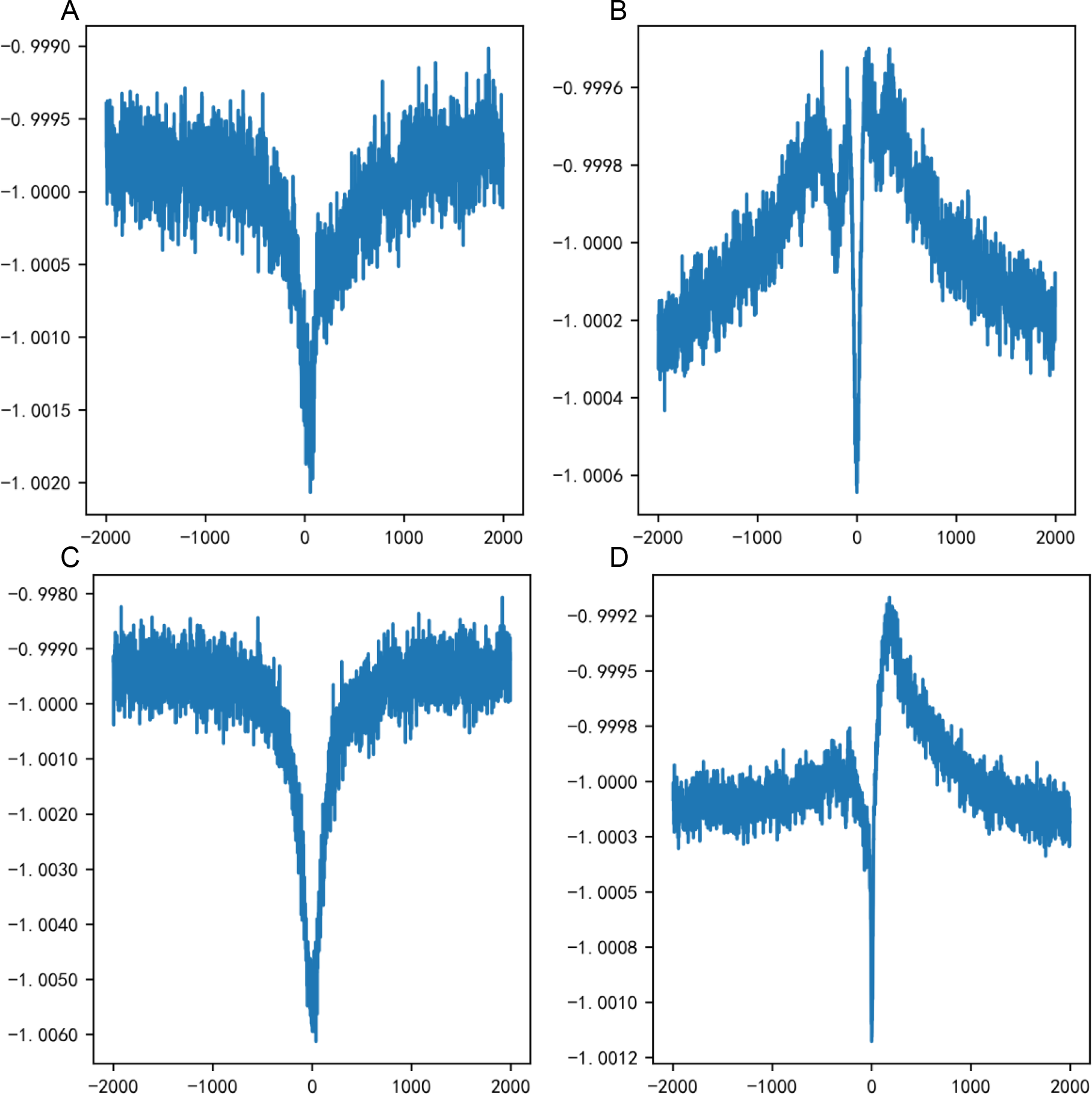
The average normalized binding energy near the TFBS. The x axis represents the distance from the TFBS. The 0 on the x-axis represents the TFBS, which is the sequence location of the highest peak defined by the ChIP-seq data. The y axis represents the averaged normalized binding energy. The binding energy is calculated using the AEEscape algorithm. (The energy used is the same as the energy in the orange boxes in Fig. 2 and Fig. 4, respectively.) (A) The binding energy around the TFBS for WRKY. (B) The binding energy around the TFBS for PU.1. (C) The binding energy around the TFBS for GR. (D) The binding energy around the TFBS for MYC2.

Since the energy funnel is derived directly from the binding energy predicted by AEEscape and the m-mer (m represents the length of the AE, e.g. for WRKY it is 3) base composition of the target sequence, the funnel must depend on the nature of the base composition around the TFBS. To investigate this, we quantify it by the average AE frequency bias b(r) adapted from (28) (Equation 8 – 9)). I(x) is an indicator function. It equals 1 if the m-mer sequence at x is the same as the examined element, and 0 otherwise. ***F***_***element***_ is the frequency of the element in the whole genome. The b(r) measures the difference between ***F***_***element***_ and the average frequency of the element at distance r from the target 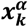. Figure 6 shows the average AE frequency bias (b(r)) for WRKY, PU.1, GR, and MYC2, respectively. There is an almost symmetrical peak around TFBS for both TFs. And the values of the peaks are the highest compared to all other m-mers for a given TF (see Fig. S12 and Fig. S14 – Fig. S16). To assess whether the AE gradient represents a unique genomic feature for TF binding, we computed the average frequency bias of all feasible 2-mer or 4-mer short sequences (Fig. S11 and Fig. S13) for WRKY. For all 2-mer short sequences, no conspicuous symmetrical peak was observed. For all 4-mer short sequences, the majority of the sequences containing the AE (GAC or GTC) exhibited a symmetrical peak. Upon comparison with the 3-mer frequency bias (Fig. S12), where the AE (GAC or GTC) displayed the highest symmetrical peak, we deduce that the AE indeed exists and exerts an in vivo role.

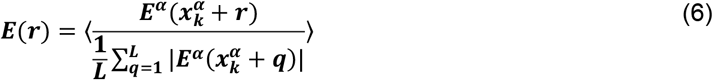

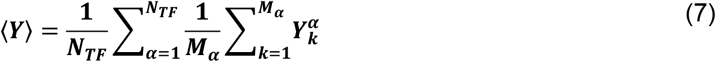

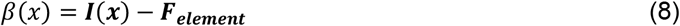

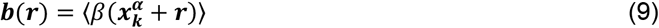

**Figure 6.**
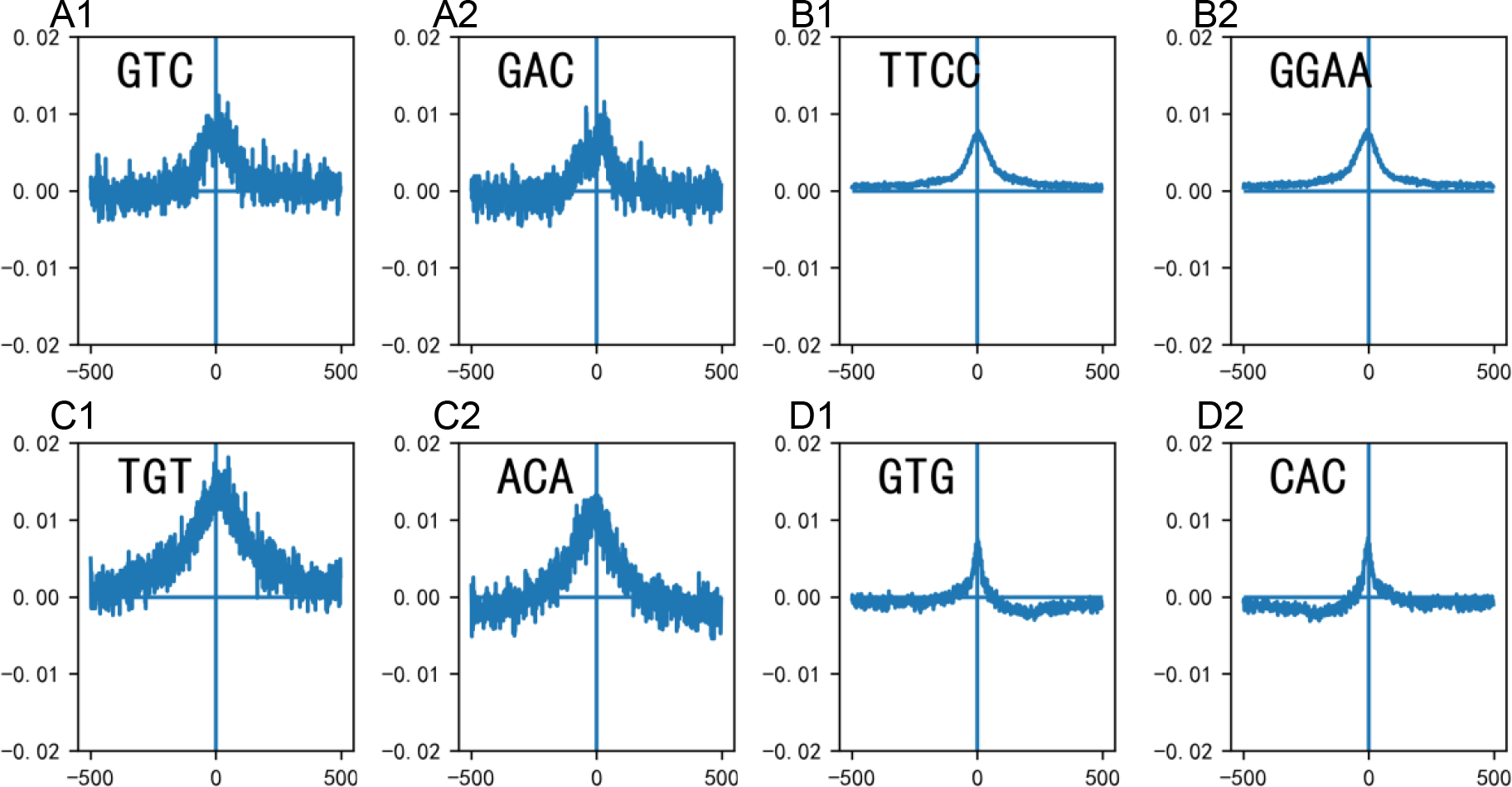
The average AE frequency bias (b(r) in the text) for different TFs in the vicinity of the TFBS. The TFBS is represented by the number zero on the x-axis. The TFBS is the sequence location of the highest peak defined by the ChIP-seq data. (A1 – A2) the average AE frequency bias around the TFBS for WRKY. (B1 – B2) the average AE frequency bias around the TFBS for PU.1. (C1 – C2) the average AE frequency bias around the TFBS for GR. (D1 – D2) the average AE frequency bias around the TFBS for MYC2.

### DISCUSSION

In this work, we detected and confirmed that there is a short m-mer sequence called AE in the binding of TF and DNA. We first did KaScape experiments of *At*WRKY1N, PU.1 DBD, cGASN, GR DBD, and MYC2 DBD. From the KaScape experiment data, we found fractal-like patterns rather than a pattern where only the motif consensus contains the lowest binding energy with all other high binding energy sequences. This implies that the AE dominates the binding energy. We further developed the AEEscape method to identify, and detect the AE and calculate its binding energy. Taking WRKY as an example, our AEEscape method shows that there is an AE “GAC” (or “GTC” in reverse complementary) for WRKY, which is consistent with the core sequence described in the paper (46,47). In their paper, the core sequence “GAC” of WRKY was determined by WRKY-dsDNA complex structure analysis and ITC (isothermal titration calorimetry) experiments (46) or by comparison between DPI-ELISA screens of WRKY family proteins (47). However, our AEEscape method extracts, defines, and validates the AE in a comprehensive and simple manner. We further analyzed the PBM data for WRKY (Fig. S17) and PU.1 (Fig. S18) on a K-mer map respectively. We found AE also exists in the PBM experiments which is another kind of independent experiment. From the AEEscape predicted maps, the binding energy of the AE is the lowest among all potential m-mers, and the difference between the AE and other m-mers is marginal. According to the structural analysis of DNA-protein complexes (Fig. S19), the AEs are situated within the major groove of B-form DNA and display the greatest degree of thermodynamic stability among the constituent nucleobases. The AE is within a widely accepted PWM motif. For example, the AE of WRKY is GAC (or GTC in complementary form) and the W-box motif of WRKY is “TTGACC/T” (48). The AEs of the PU.1, GR, and MYC2 DBDs are all located within the high-information PWM motif. (Compare Fig. S24 and the AEs for the corresponding TF DBDs).

As mentioned above, the AEEscape method is based on BEESEM. To compare the accuracy and interpretability of AEEscape and BEESEM, we used the input-output dataset for *At*WRKY1N (the same dataset in the KaScape experimental paper (11)) as a benchmark dataset. Figure S20 shows the predicted binding energy by AEEscape for all 3-mer sequences at each location in the random base region for R1N4, R1N5, R1N6, and R1N7 KaScape experimental data, respectively. The binding energy of GAC is lowest at the first site (first column) and tends to be relatively low at other sites as well. The binding energy of GTC is also quite low in some sub-figures, such as Fig. S20 A2 and Fig. S20 C3. Almost all sub-figures have a uniform background pattern (e.g., the binding energy of the AC diagonal is relatively low), which is due to noise. These results are consistent with Fig. 2B1 – Fig. 2B3. The training process converges when it stops at the 10th EM loop (Table S3). We then use the BEESEM algorithm to predict the PWM parameters for the same data set (Fig. S21). The EM loop number of the BEESEM algorithm is 10 and the result is convergent (Table S3). When the random sequence type is R1N4 or R1N6 in the KaScape experiment dataset, the PWM contains GTC. However, for other random sequence-type KaScape experiments, it is predicted to be AT-rich. Figure S22 and Figure S23 show the training efficiency comparison between BEESEM and AEEscape algorithms. The AEEscape method outperforms the BEESEM method. The AEEscape method can detect AEs more sensitively and derive the binding energy landscape more accurately than the BEESEM algorithm. The AEEscape method can help us to better understand the binding mechanism between TFs and dsDNA.

We have revealed an energetic funnel around TFBS through the ChIP-seq datasets (42-45) as well as the m-mer energy landscape derived from the AEEscape method. However, when we use the PWM motif (obtained from the Jaspar website Fig. S24) to calculate the binding energy around TFBS, the energetic funnel disappears or is not obvious for most proteins (Fig. S25). The binding energy funnel is revealed by the m-mer energy landscape derived from AEEscape, but not by the PWM motif because the AE is shorter than the PWM motif and the probability of occurrence of an AE density gradient in the sequences around the TFBS is greater. The numerical simulations in paper (28) already show that the energy funnel can significantly increase the probability of finding TFBSs, which facilitates facilitated diffusion. Thus, the existence of an AE density gradient around TFBS will facilitate TFs to find their targets faster than diffusion-limited rates. The AE is the core in the widely known motif that is sufficient for TF binding. This means that the AE can enhance the binding of TF and dsDNA, which can help the TF to interact with dsDNA in one dimension and make the target finding reliable. The short length of the AE also indicates that the energy difference between the AE and other elements is relatively small. This small free energy barrier will not make the binding too strong, and the small free energy barrier will facilitate turnover and not trap TF for a long time.

By examining sequences flanking TFBSs identified from the ChIP-seq dataset, we discovered that there is a gradient of AEs density around TFBSs. Although it has been suggested that the flanking sequence could influence the binding of TFs to their target, it was previously thought that this was due to dsDNA properties such as flexibility, stability, and minor groove width (49) or non-specific binding (e.g., AT-rich sequence (28)); it did not take into account the correlation between the flanking sequence and the specific binding site. Here we show that the flanking sequence is also related to the specific binding sites by the AE density gradient. The AE density gradient also suggests that the existence of TF around TFBS is a kind of probability. It breaks the convention that the TF recognizes its target in a 0 or 1 way (it needs to recognize a consensus motif such as the W-box, and search it on the genomic sequence to determine the TFBS) (50-53). The probability of binding to the TFBS increases along the flanking dsDNA and is highest at the TFBS. As long as there are enough buffering TFs around the TFBS, it will help to promote or block the transcription machinery and also help to form a combinatorial transcription complex. Note that there are other m-mer density gradients around TFBS besides the AE (Fig. S11 – Fig. S14). This may help to further localize the TF, maybe another AE sequence, or perhaps the AEs of other TFs, considering the combinatorial interaction among TFs that are ubiquitous in the eukaryotic transcription (54,55).

The new search mechanism proposed here establishes the molecular aspect of the interaction between TF and flanking DNA regions around TFBS. It also clarifies the role of the regulatory region around the TFBS and further indicates that the AE density gradient in the regulatory region is important. The mechanism implies that rapid and accurate TF recognition depends not only on the TF itself but also on the flanking sequence composition (the AE density gradient) around the TFBS. Thus, there is a co-evolution between the flanking sequence around the TFBS and the TF. This is because the density along the dsDNA is essential rather than a critical specific site. The mechanism is resistant to mutation and can withstand a greater burden of mutations in the flanking sequence around the TFBS (56).

According to the mechanism, a single molecule experiment can be designed to test whether it supports facilitated diffusion. The mechanism provides a new strategy for synthetic promoter design. Calculating the density of elements across the genome can help determine the AE for a particular TF and hopefully predict the potentially regulated genes. The TF binding used in the paper is a one-site binding (WRKY, PU.1) or two-site binding (GR, MYC2), and the dsDNA used in the KaScape experiments are free of histones. However, the eukaryotic genome is nucleosomal DNA (57).

Whether the mechanism applies to the interaction between nucleosomal dsDNA and pioneer factor remains to be investigated (58). Further research is also needed on TFs that require the combination of other TFs to function (59,60). The approach described in this study can be extended to address questions that are more relevant to the actual eukaryotic genome.

## AVAILABILITY

AEEscape is available from the GitHub repository (https://github.com/NinYuan/AEEscape.git)

## SUPPLEMENTARY DATA

Supplementary Data are available at NAR online.

## ACKNOWLEDGEMENT

We thank Wenhao Wu for his help with the KaScape experiment.

## REFERENCES

1. Roeder, R.G. (2019) 50+ years of eukaryotic transcription: an expanding universe of factors and mechanisms. Nat Struct Mol Biol., 26, 783–791.

2. Latchman, D.S. (1993) Transcription factors: an overview. Int J Exp Pathol., 74, 417–422.

3. Hardison, R.C. and Taylor, J. (2012) Genomic approaches towards finding cis-regulatory modules in animals. Nat Rev Genet., 13, 469–483.

4. Stormo, G.D., Schneider, T.D., Gold, L. and Ehrenfeucht, A. (1982) Use of the ‘Perceptron’ algorithm to distinguish translational initiation sites in E. coli. Nucleic Acids Res., 10, 2997–3011.

5. Stormo, G.D. (2013) Modeling the specificity of protein-DNA interactions. Quant Biol., 1, 115–130.

6. Stormo, G.D. (2015) DNA Motif Databases and Their Uses. Curr Protoc Bioinformatics., 51, 2.15.11–12.15.16.

7. Jolma, A., Kivioja, T., Toivonen, J., Cheng, L., Wei, G., Enge, M., Taipale, M., Vaquerizas, J.M., Yan, J., Sillanpää, M.J. et al. (2010) Multiplexed massively parallel SELEX for characterization of human transcription factor binding specificities. Genome Res., 20, 861–873.

8. Berger, M.F., Philippakis, A.A., Qureshi, A.M., He, F.S., Estep, P.W. and Bulyk, M.L. (2006) Compact, universal DNA microarrays to comprehensively determine transcription-factor binding site specificities. Nat Biotechnol., 24, 1429–1435.

9. Maerkl, S.J. and Quake, S.R. (2007) A systems approach to measuring the binding energy landscapes of transcription factors. Science, 315, 233–237.

10. Rockel, S., Geertz, M. and Maerkl, S.J. (2012) MITOMI: a microfluidic platform for in vitro characterization of transcription factor-DNA interaction. Methods Mol Biol., 786, 97–114.

11. Chen, H., Xu, Y., Jin, J. and Su, X.-d. (2023) KaScape: a sequencing-based method for global characterization of protein-DNA binding affinity. Sci Rep., 13, 16595.

12. Johnson, D.S., Mortazavi, A., Myers, R.M. and Wold, B. (2007) Genome-wide mapping of in vivo protein-DNA interactions. Science, 316, 1497–1502.

13. Bulyk, M.L. (2003) Computational prediction of transcription-factor binding site locations. Genome Biol., 5, 201.

14. Stormo, G.D. (2000) DNA binding sites: representation and discovery. Bioinformatics, 16, 16–23.

15. Wasserman, W.W. and Sandelin, A. (2004) Applied bioinformatics for the identification of regulatory elements. Nat Rev Genet., 5, 276–287.

16. GuhaThakurta, D. (2006) Computational identification of transcriptional regulatory elements in DNA sequence. Nucleic Acids Res., 34, 3585–3598.

17. Jones, S.J. (2006) Prediction of genomic functional elements. Annu Rev Genomics Hum Genet., 7, 315–338.

18. Pollard, D.A., Bergman, C.M., Stoye, J., Celniker, S.E. and Eisen, M.B. (2004) Benchmarking tools for the alignment of functional noncoding DNA. BMC Bioinformatics, 5, 6.

19. Gershenzon, N.I., Stormo, G.D. and Ioshikhes, I.P. (2005) Computational technique for improvement of the position-weight matrices for the DNA/protein binding sites. Nucleic Acids Res., 33, 2290–2301.

20. O’Flanagan, R.A., Paillard, G., Lavery, R. and Sengupta, A.M. (2005) Non-additivity in protein-DNA binding. Bioinformatics, 21, 2254–2263.

21. Barash, Y., Elidan, G., Friedman, N. and Kaplan, T. (2003), Proceedings of the seventh annual international conference on Research in computational molecular biology. Association for Computing Machinery, Berlin, Germany, pp. 28–37.

22. Zhang, Q., Shen, Z. and Huang, D.-S. (2019) Modeling in-vivo protein-DNA binding by combining multiple-instance learning with a hybrid deep neural network. Sci Rep., 9, 8484.

23. Jana, T., Brodsky, S. and Barkai, N. (2021) Speed-Specificity Trade-Offs in the Transcription Factors Search for Their Genomic Binding Sites. Trends Genet., 37, 421–432.

24. Riggs, A.D., Bourgeois, S. and Cohn, M. (1970) The lac repressor-operator interaction. 3. Kinetic studies. J Mol Biol., 53, 401–417.

25. Berg, O.G., Winter, R.B. and von Hippel, P.H. (1981) Diffusion-driven mechanisms of protein translocation on nucleic acids. 1. Models and theory. Biochemistry, 20, 6929–6948.

26. Berg, O.G., Winter, R.B. and von Hippel, P.H. (1982) How do genome-regulatory proteins locate their DNA target sites? Trends Biochem Sci., 7, 52–55.

27. von Hippel, P.H. and Berg, O.G. (1989) Facilitated target location in biological systems. J Biol Chem., 264, 675–678.

28. Cencini, M. and Pigolotti, S. (2018) Energetic funnel facilitates facilitated diffusion. Nucleic Acids Res., 46, 558–567.

29. Wunderlich, Z. and Mirny, L.A. (2009) Different gene regulation strategies revealed by analysis of binding motifs. Trends Genet., 25, 434–440.

30. Cohen, D.M., Lim, H.W., Won, K.J. and Steger, D.J. (2018) Shared nucleotide flanks confer transcriptional competency to bZip core motifs. Nucleic Acids Res., 46, 8371–8384.

31. Shahein, A., López-Malo, M., Istomin, I., Olson, E.J., Cheng, S. and Maerkl, S.J. (2022) Systematic analysis of low-affinity transcription factor binding site clusters in vitro and in vivo establishes their functional relevance. Nat Commun., 13, 5273.

32. Siggia, E.D. (2005) Computational methods for transcriptional regulation. Curr Opin Genet Dev., 15, 214–221.

33. Castellanos, M., Mothi, N. and Muñoz, V. (2020) Eukaryotic transcription factors can track and control their target genes using DNA antennas. Nat Commun., 11, 540.

34. Brackley, C.A., Cates, M.E. and Marenduzzo, D. (2012) Facilitated diffusion on mobile DNA: configurational traps and sequence heterogeneity. Phys Rev Lett., 109, 168103.

35. Horton, C.A., Alexandari, A.M., Hayes, M.G.B., Marklund, E., Schaepe, J.M., Aditham, A.K., Shah, N., Suzuki, P.H., Shrikumar, A., Afek, A. et al. (2023) Short tandem repeats bind transcription factors to tune eukaryotic gene expression. Science, 381, eadd1250.

36. Sharma, D., Sharma, K., Mishra, A., Siwach, P., Mittal, A. and Jayaram, B. (2023) Molecular dynamics simulation-based trinucleotide and tetranucleotide level structural and energy characterization of the functional units of genomic DNA. Phys Chem Chem Phys., 25, 7323–7337.

37. Ruan, S., Swamidass, S.J. and Stormo, G.D. (2017) BEESEM: estimation of binding energy models using HT-SELEX data. Bioinformatics, 33, 2288–2295.

38. Benos, P.V., Bulyk, M.L. and Stormo, G.D. (2002) Additivity in protein-DNA interactions: how good an approximation is it? Nucleic Acids Res., 30, 4442–4451.

39. Roulet, E., Busso, S., Camargo, A.A., Simpson, A.J., Mermod, N. and Bucher, P. (2002) High-throughput SELEX SAGE method for quantitative modeling of transcription-factor binding sites. Nat Biotechnol., 20, 831–835.

40. Jolma, A., Yan, J., Whitington, T., Toivonen, J., Nitta, K.R., Rastas, P., Morgunova, E., Enge, M., Taipale, M., Wei, G. et al. (2013) DNA-binding specificities of human transcription factors. Cell, 152, 327–339.

41. Hao, B.-l., Lee, H.C. and Zhang, S.-y. (2000) Fractals related to long DNA sequences and complete genomes. Chaos, Solitons & Fractals, 11, 825–836.

42. Birkenbihl, R.P., Kracher, B., Roccaro, M. and Somssich, I.E. (2017) Induced Genome-Wide Binding of Three Arabidopsis WRKY Transcription Factors during Early MAMP-Triggered Immunity. Plant Cell, 29, 20–38.

43. Gertz, J., Savic, D., Varley Katherine E., Partridge, E.C., Safi, A., Jain, P., Cooper Gregory M., Reddy Timothy E., Crawford Gregory E. and Myers Richard M. (2013) Distinct Properties of Cell-Type-Specific and Shared Transcription Factor Binding Sites. Mol Cell., 52, 25–36.

44. Greulich, F., Wierer, M., Mechtidou, A., Gonzalez-Garcia, O. and Uhlenhaut, N.H. (2021) The glucocorticoid receptor recruits the COMPASS complex to regulate inflammatory transcription at macrophage enhancers. Cell Rep., 34, 108742.

45. Zander, M., Lewsey, M.G., Clark, N.M., Yin, L., Bartlett, A., Saldierna Guzmán, J.P., Hann, E., Langford, A.E., Jow, B., Wise, A. et al. (2020) Integrated multi-omics framework of the plant response to jasmonic acid. Nat Plants., 6, 290–302.

46. Xu, Y.P., Xu, H., Wang, B. and Su, X.D. (2020) Crystal structures of N-terminal WRKY transcription factors and DNA complexes. Protein Cell, 11, 208–213.

47. Brand, L.H., Fischer, N.M., Harter, K., Kohlbacher, O. and Wanke, D. (2013) Elucidating the evolutionary conserved DNA-binding specificities of WRKY transcription factors by molecular dynamics and in vitro binding assays. Nucleic Acids Res., 41, 9764–9778.

48. Ciolkowski, I., Wanke, D., Birkenbihl, R.P. and Somssich, I.E. (2008) Studies on DNA-binding selectivity of WRKY transcription factors lend structural clues into WRKY-domain function. Plant Mol Biol., 68, 81–92.

49. Yella, V.R., Bhimsaria, D., Ghoshdastidar, D., Rodríguez-Martínez, J.A., Ansari, A.Z. and Bansal, M. (2018) Flexibility and structure of flanking DNA impact transcription factor affinity for its core motif. Nucleic Acids Res., 46, 11883–11897.

50. Krystel, J. and Ayyanathan, K. (2012) An efficient and cost-effective protocol for selecting transcription factor binding sites that reduces isotope usage. J Biomol Tech., 23, 40–46.

51. Lee, C. and Huang, C.H. (2012) Searching for transcription factor binding sites in vector spaces. BMC Bioinformatics, 13, 215.

52. Bryne, J.C., Valen, E., Tang, M.H., Marstrand, T., Winther, O., da Piedade, I., Krogh, A., Lenhard, B. and Sandelin, A. (2008) JASPAR, the open access database of transcription factor-binding profiles: new content and tools in the 2008 update. Nucleic Acids Res., 36, D102–106.

53. Hashim, F.A., Mabrouk, M.S. and Al-Atabany, W. (2019) Review of Different Sequence Motif Finding Algorithms. Avicenna J Med Biotechnol., 11, 130–148.

54. Ravasi, T., Suzuki, H., Cannistraci, C.V., Katayama, S., Bajic, V.B., Tan, K., Akalin, A., Schmeier, S., Kanamori-Katayama, M., Bertin, N. et al. (2010) An atlas of combinatorial transcriptional regulation in mouse and man. Cell, 140, 744–752.

55. Morgunova, E. and Taipale, J. (2017) Structural perspective of cooperative transcription factor binding. Curr Opin Struct Biol., 47, 1–8.

56. Crocker, J., Abe, N., Rinaldi, L., McGregor, A.P., Frankel, N., Wang, S., Alsawadi, A., Valenti, P., Plaza, S., Payre, F. et al. (2015) Low affinity binding site clusters confer hox specificity and regulatory robustness. Cell, 160, 191–203.

57. Michael, A.K. and Thomä, N.H. (2021) Reading the chromatinized genome. Cell, 184, 3599–3611.

58. Zhu, F., Farnung, L., Kaasinen, E., Sahu, B., Yin, Y., Wei, B., Dodonova, S.O., Nitta, K.R., Morgunova, E., Taipale, M. et al. (2018) The interaction landscape between transcription factors and the nucleosome. Nature, 562, 76–81.

59. Reiter, F., Wienerroither, S. and Stark, A. (2017) Combinatorial function of transcription factors and cofactors. Curr Opin Genet Dev., 43, 73–81.

60. Panne, D., Maniatis, T. and Harrison, S.C. (2007) An atomic model of the interferon-beta enhanceosome. Cell, 129, 1111–1123.

